# ASaiM: a Galaxy-based framework to analyze raw shotgun data from microbiota

**DOI:** 10.1101/183970

**Authors:** Bérénice Batut, Kévin Gravouil, Clémence Defois, Saskia Hiltemann, Jean-François Brugère, Eric Peyretaillade, Pierre Peyret

## Abstract

**Background:** New generation of sequencing platforms coupled to numerous bioinformatics tools has led to rapid technological progress in metagenomics and metatranscriptomics to investigate complex microorganism communities. Nevertheless, a combination of different bioinformatic tools remains necessary to draw conclusions out of microbiota studies. Modular and user-friendly tools would greatly improve such studies.

**Findings:** We therefore developed ASaiM, an Open-Source Galaxy-based framework dedicated to microbiota data analyses. ASaiM provides a curated collection of tools to explore and visualize taxonomic and functional information from raw amplicon, metagenomic or metatranscriptomic sequences. To guide different analyses, several customizable workflows are included. All workflows are supported by tutorials and Galaxy interactive tours to guide the users through the analyses step by step. ASaiM is implemented as Galaxy Docker flavour. It is scalable to many thousand datasets, but also can be used a normal PC. The associated source code is available under Apache 2 license at https://github.com/ASaiM/framework and documentation can be found online (http://asaim.readthedocs.io/)

**Conclusions:** Based on the Galaxy framework, ASaiM offers sophisticated analyses to scientists without command-line knowledge. ASaiM provides a powerful framework to easily and quickly explore microbiota data in a reproducible and transparent environment.

## Findings

### Background

The study of microbiota and microbial communities has been facilitated by the evolution of sequencing techniques and the development of metagenomics and metatranscriptomics. These techniques are giving insight into phylogenetic properties and metabolic components of microbial communities. However, meta’omic data exploitation is not trivial due to the large amount of data, high variability, incompleteness of reference databases, difficulty to find, configure, use and combine the dedicated bioinformatics tools, etc. Hence, to extract useful information, a sequenced microbiota sample has to be processed by sophisticated workflows with numerous successive bioinformatics steps [1]. Each step may require execution of several tools or software programs. For example, to extract taxonomic information with the widely used QIIME [2] or Mothur [[3], at least 10 different tools with at least 4 parameters each are needed. Designed for amplicon data, both QIIME and Mothur can not be directly applied to shotgun metagenomics data. In addition, the tools can be complex to use; they are command-line tools and may require computational resources specially for the metagenomics datasets. In this context, selecting the best tools, configuring them to use the correct parameters and appropriate computational resources and combining them together in an analysis chain is a complex and error-prone process. These issues and the involved complexity are blocking scientist from participating in the analysis of their own data. Furthermore, bioinformatics tools are often manually executed and/or patched together with custom scripts. These practices raise doubts about a science gold standard: reproducibility [3,4]. Web services and automated pipelines such as MG-RAST [5] and EBI metagenomics [6] offer solutions to the accessibility issue. However, these web services work as a black box and are lacking in transparency, flexibility and even reproducibility as the version and parameters of the tools are not always available. Alternative approaches to improve accessibility, modularity and reproducibility can be found in open-source workflow systems such as Galaxy [6–8]. Galaxy is a lightweight environment providing a web-based, intuitive and accessible user interface to command-line tools, while automatically managing computation and transparently managing data provenance and workflow scheduling [6–8]. More than 4,500 tools can be used inside Galaxy environments. The tools can be selected and combined to build Galaxy flavors focusing on specific type of analysis, *e.g.* the Galaxy RNA workbench [9].

In this context, we developed ASaiM (Auvergne Sequence analysis of intestinal Microbiota), an Open-Source opinionated Galaxy-based framework. It integrates tools and workflows dedicated to microbiota analyses with an extensive documentation (http://asaim.readthedocs.org) and training support.

### Goals of ASaiM

ASaiM is developed as a modular, accessible, redistributable, sharable and user-friendly framework for scientists working with microbiota data. This framework is unique in combining curated tools and workflows and providing easy access for scientists.

ASaiM is based on four pillars: 1) easy and stable dissemination via Galaxy, Docker and conda, 2) a comprehensive set of metagenomics related tools, 3) a set of predefined and tested workflows, and 4) extensive documentation and training to help scientists in their analyses.

### A framework built on the shoulders of giants

The ASaiM framework is built on existing tools and infrastructures and combine all their forces to build an easily accessible and reproducible analysis platform.

ASaiM is implemented as portable virtualized container based on Galaxy framework [8]. Galaxy provides researchers with means to reproduce their own workflows analyses, rerun entire pipelines, or publish and share them with others. Based on Galaxy, ASaiM is scalable from single CPU installations to large multi-node high performance computing environments. Deployments can be archived by using a pre-built ASaiM Docker image, which is based on the Galaxy Docker project (http://bgruening.github.io/docker-galaxy-stable) or by installing all needed components into an already existing Galaxy instance. This ASaiM Docker instance is customized with a variety of selected tools, workflows, Interactive tours and data that have been added as additional layers on top of the generic Galaxy Docker instance. The containerization keeps the deployment task to a minimum. The selected Galaxy tools are automatically installed from the Galaxy ToolShed [10] (https://toolshed.g2.bx.psu.edu/) using the Galaxy API BioBlend [11] and the installation of the tools and their dependencies are automatically resolved using packages available through Bioconda (https://bioconda.github.io). We migrated then 10 tools/suites of tools and their dependencies to Bioconda (*e.g.* HUMAnN2) and integrated 14 suites into Galaxy (*e.g.*QIIME with around forty tools).

The containerization as well as the packaging with conda enables automatic continuous integration tests at different levels: dependencies (BioConda), tool integration in Galaxy, Galaxy itself and at ASaiM level. Together with strict version management on all levels, this contributes to a high degree of error-control and reproducibility.

### Tools for microbiota data analyses

The tools integrated in ASaiM can be seen in Table 1. They are expertly selected for their relevance with regard to microbiota studies, such as Mothur [3], QIIME [2], MetaPhlAn2 [12], HUMAnN2 [13] or tools used in existing pipelines such as EBI Metagenomics’ one. We also added general tools used in sequence analysis such as quality control, mapping or similarity search tools.

**Table 1:**
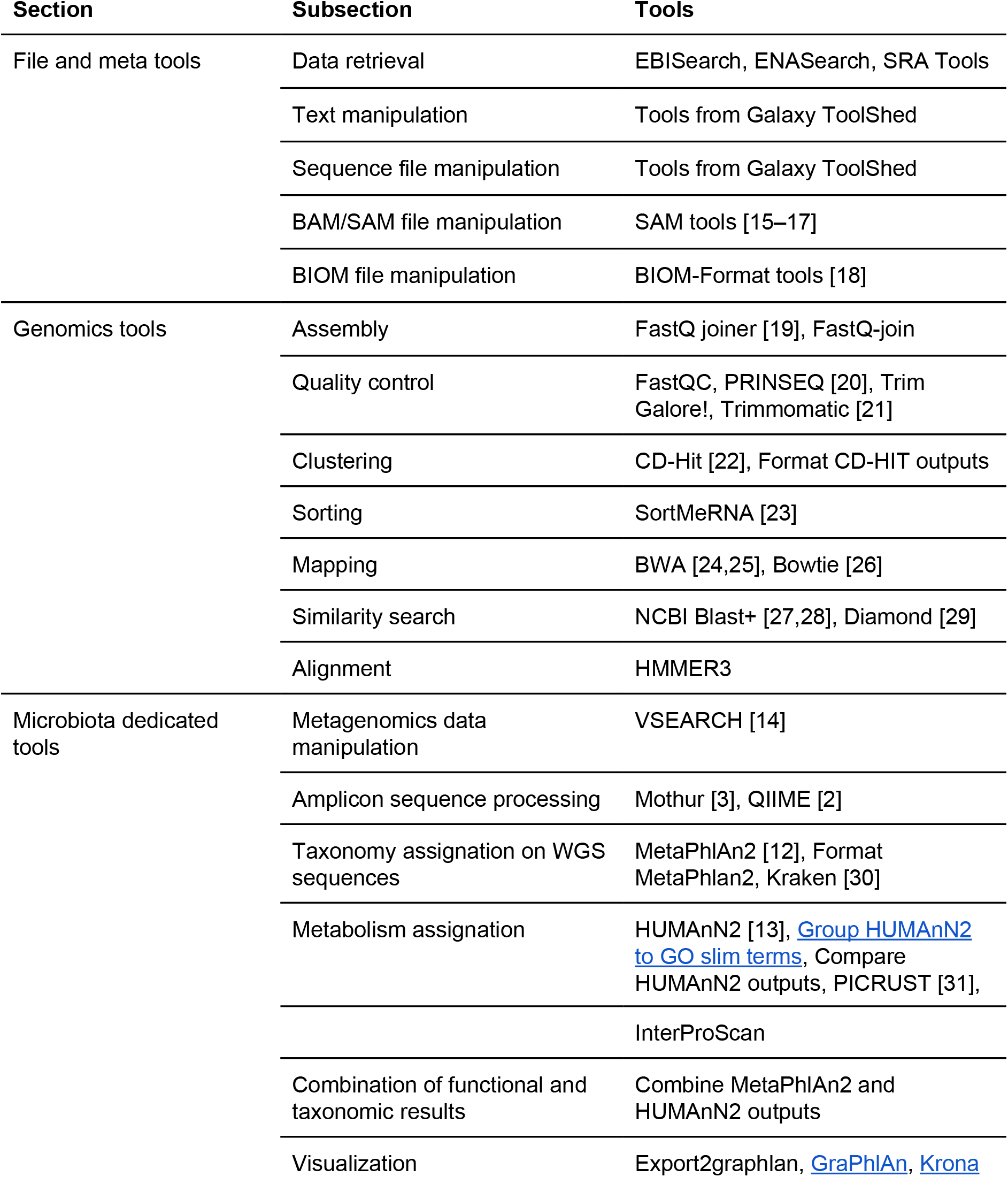
Available tools in ASaiM

This table presents the tools, organized in section and subsections to help users. A more detailed table of the available tools and some documentation can be found in the online documentation (http://asaim.readthedocs.org/)

An effort in development was made to integrate these tools into Conda and the Galaxy environment, with the help and support of the Galaxy community. We also developed two new tools to search and get data from EBI Metagenomics and ENA databases using the API of the databases (EBISearch and ENASearch) and a tool to group HUMAnN2 outputs into Gene Ontology Slim Terms. Tools inside ASaiM are organized to make them findable and documented (http://asaim.readthedocs.org/).

#### Diverse source of data

Any easy way to upload user-data into ASaiM is provided by an web-interface or more sophisticated via FTP or SFTP. Moreover, we added specialised tools that can interact with external databases like NCBI, ENA or EBI Metagenomics to query them and download data into the framework.

#### Visualization of the data

An analysis often ends with summarizing figures that conclude and represent the findings. ASaiM includes standard interactive plotting tools to draw bar charts and scatter plots from all kinds of tabular data. Phinch visualization is also included to interactively visualize and explore any BIOM file, and generate different types of ready-to-publish figures. We also integrated two other tools to explore and represent the community structure from outputs of MetaPhlAn: KRONA (Ondov et al. 2011)and GraPhlAn. Moreover, as in any Galaxy instance, other visualization are included such Phyloviz for phylogenetic trees or the Genome browser Trackster for visualizing SAM/BAM, BED, GFF/GTF, WIG, bigWig, bigBed, bedGraph, and VCF datasets.

### Workflows

Each tool can be used separately in an explorative manner or multiple tools can be orchestrated inside workflows passing raw data to the data reduction step, to information extraction and visualization. To assist in microbiota analyses, several default but customizable workflows are proposed in ASaiM. All the available workflows with tool and parameter choices are documented (http://asaim.readthedocs.org/).

#### Analysis of raw metagenomic or metatranscriptomic shotgun data

A workflow quickly produces, from raw metagenomic or metatranscriptomic shotgun data, accurate and precise taxonomic assignations, wide extended functional results and taxonomically related metabolism information (Figure 1). This workflow consists of i) processing with quality control/trimming (FastQC and Trim Galore!) and dereplication (VSearch [14]; ii) taxonomic analyses with assignation (MetaPhlAn2 [12]) and visualization (KRONA, GraPhlAn); iii) functional analyses with metabolic assignation and pathway reconstruction (HUMAnN2 [13]); iv) functional and taxonomic combination with developed tools combining HUMAnN2 and MetaPhlAn2 outputs.

This workflow has been tested on two mock metagenomic datasets with controlled communities (Supplementary material). We have compared the extracted taxonomic and functional information to such information extracted with the EBI metagenomics’ pipeline and to the expectations from the mock datasets. With ASaiM, we generate more accurate and precise data for taxonomic analyses (Figure 2): we can access information at the level of the species. More informative data for metabolic description (gene families, gene ontologies, pathways, etc) are also extracted with ASaiM compared to the ones available on EBI metagenomics. With this workflow, we can investigate which taxons are involved in a specific pathway or a gene family (*e.g.* involved species and their relative involvement in different step of fatty acid biosynthesis pathways, Figure 3).

**Figure 1:**
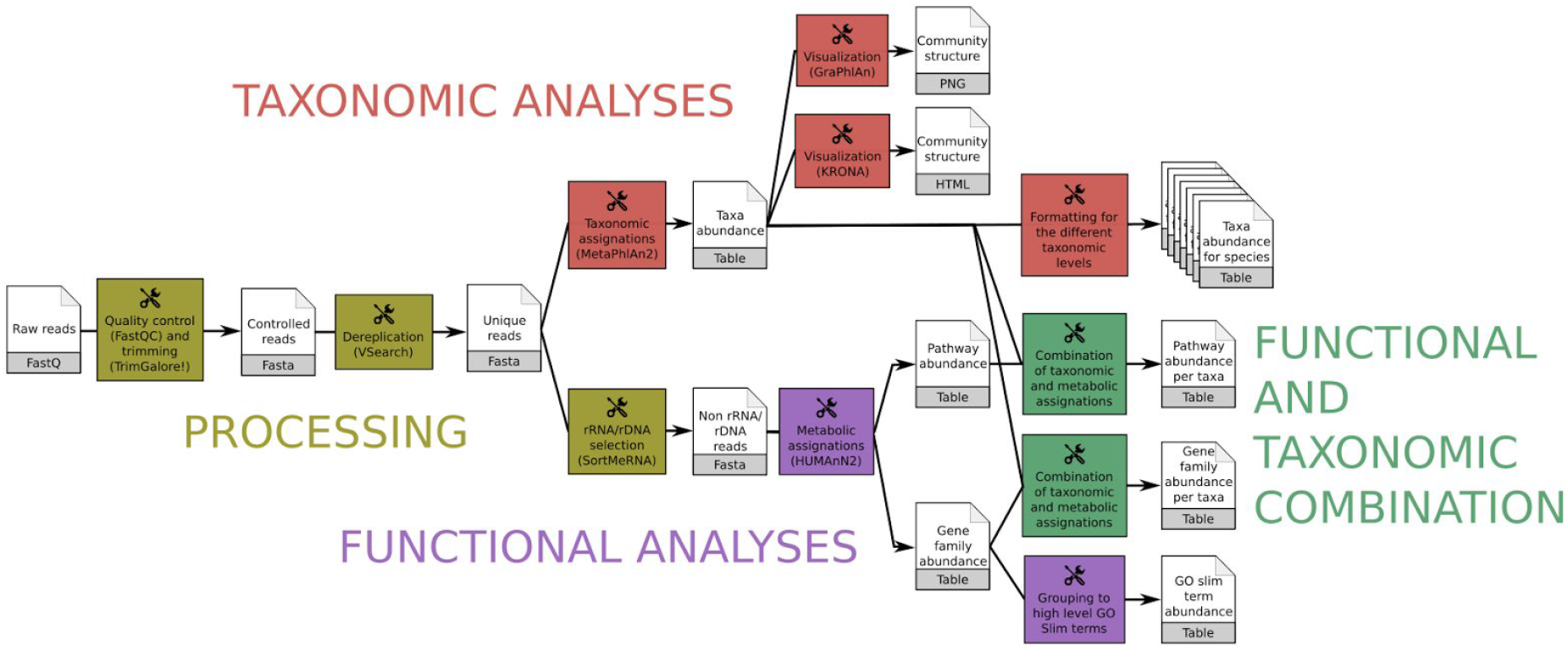
Main ASaiM workflow to analyze raw sequences. This workflow takes as input a dataset of raw shotgun sequences (in FastQ format) from microbiota, preprocess it (yellow boxes), extracts taxonomic (red boxes) and functional (purple boxes) assignations and combines them (green boxes). Image available under CC-BY license (https://doi.org/10.6084/m9.figshare.5371396.v3)

**Figure 2:**
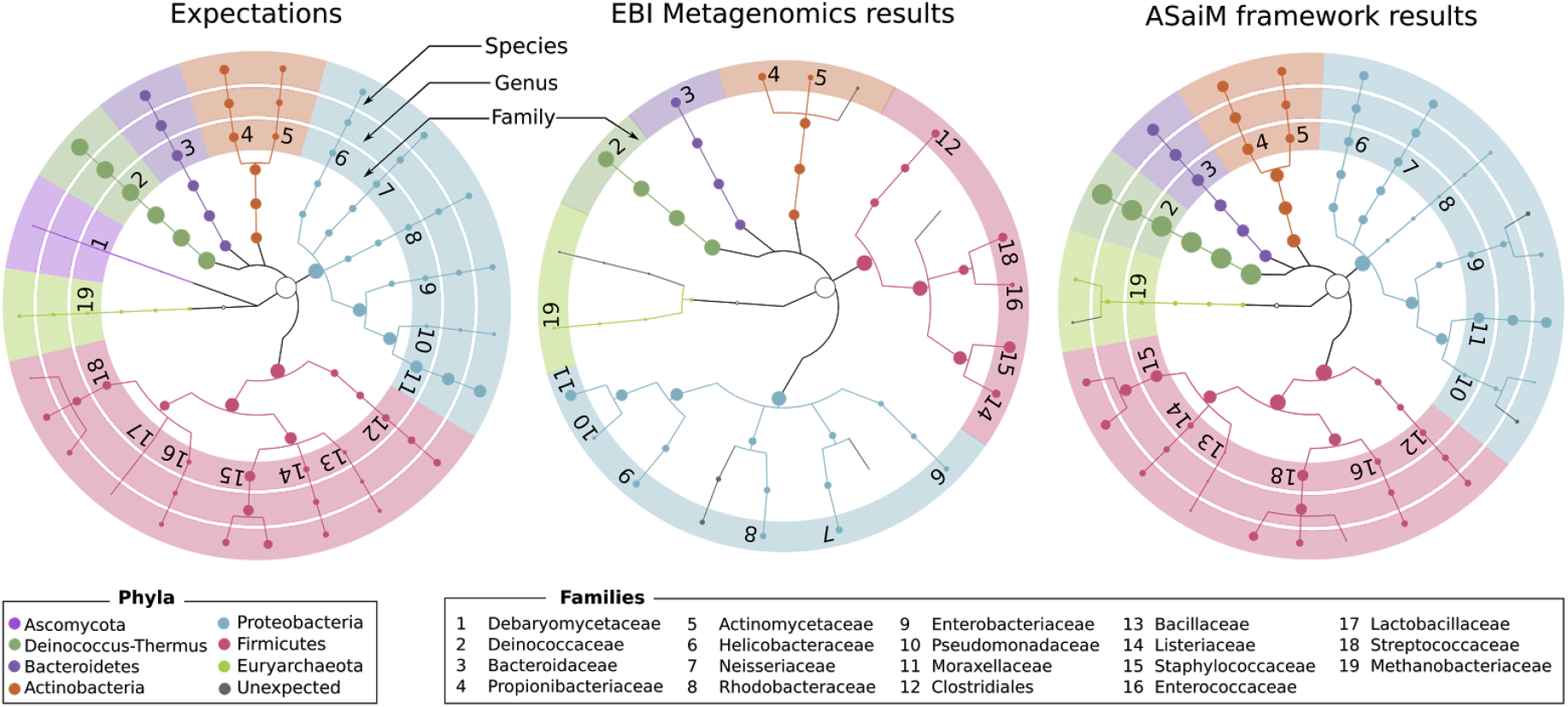
Comparisons of the community structure for SRR072233. This figure compares the community structure between the expectations (mapping of the sequences on the expected genomes), data found on EBI Metagenomics database (extracted with the EBI Metagenomics pipeline) and the results of the main ASaiM workflow (Figure 1).

**Figure 3:**
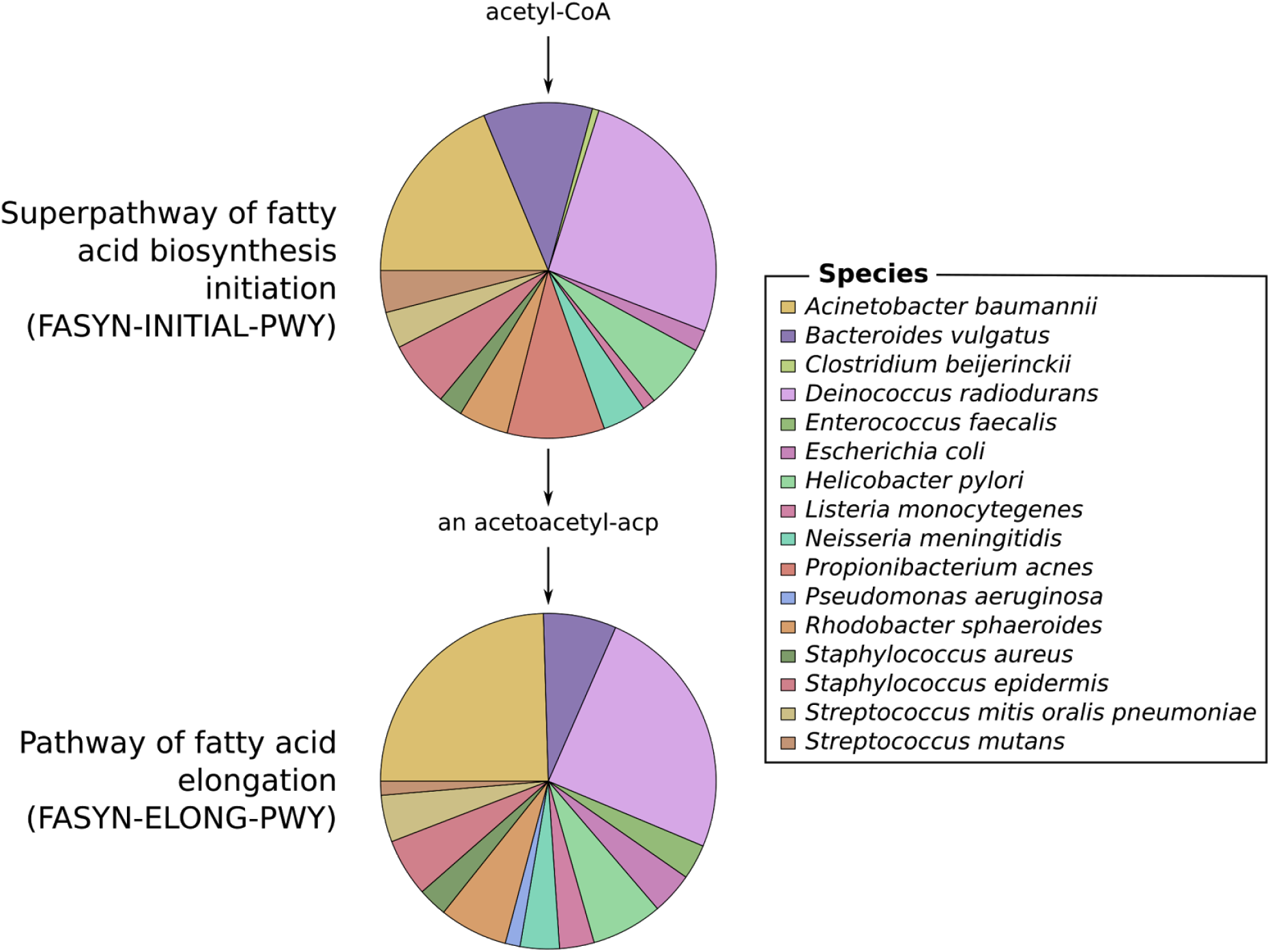
Example of an investigation of the relation between community structure and functions. The involved species and their relative involvement in fatty acid biosynthesis pathways have been extracted with ASaiM workflow (Figure 1) for SRR072233

For the tests, ASaiM was deployed on a computer with Debian GNU/Linux System, 8 cores Intel(R) Xeon(R) at 2.40 GHz and 32 Go of RAM. The workflow processed the 1,225,169 and 1,386,198 454 GS FLX Titanium reads of each datasets in 4h44 and 5h22 respectively, with a stable memory usage (Supplementary material). With this workflow, it is then easy and quick to process raw microbiota data and extract diverse useful information.

#### Analysis of amplicon data

To analyze amplicon data, the Mothur and QIIME tool suites are available to ASaiM. We integrated the workflows described in tutorials of Mothur and QIIME websites, as example of amplicon data analyses as well as support for the training material. These workflows, as any workflows available in ASaiM, can be adapted for a specific analysis or used as subworkflows by the users.

#### Running as in EBI metagenomics

The tools used in the EBI Metagenomics pipeline are also available in ASaiM. We integrate then also a workflow with the same steps as the EBI Metagenomics pipeline. Analyses made in EBI Metagenomics website can be then reproduced locally, without having to wait for availability of EBI Metagenomics or to upload any data on EBI Metagenomics. However the parameters must be defined by the user as we can not find them on EBI Metagenomics documentation.

### Documentation and training

A tool or software is easier to use if it is well documented. Hence extensive documentation helps the users to be familiar with the tool and also prevents mis-usage. For ASaiM, we developed an extensive online documentation (http://asaim.readthedocs.io/), mainly to explain how to use it, how to deploy it, which tools are integrated with small documentation about these tools, which workflows are integrated and how to use them.

In addition to this online documentation, Galaxy Interactive Tours are included inside the Galaxy instance. Such tours guide users through an entire analysis in an interactive (step-by-step) way. Some tours, included in every Galaxy instance, explains how to use Galaxy. We also developed such tours dedicated specifically to the ASaiM workflows.

These interactive tours are used to complement tutorials and trainings. Some tutorials about the integrated workflows have been developed to explain step-by-step the workflows with small example datasets. Hosted in the Galaxy Training Network (GTN) GitHub repository (https://github.com/galaxyproject/training-material), the tutorials are available online at http://training.galaxyproiect.org. They have been used during several workshops on metagenomics data analysis with ASaiM as training support. These tutorials are also accessible directly from ASaiM and its documentation for self-training.

### Installation and running ASaiM

Running the containerized ASaiM simply requires to install Docker and to start the ASaiM image with:

> $ docker run ‒d ‒p 8080:80 quay.io/bebatut/asaim

Thanks to Docker, ASaiM can be installed under every operating systems, even with a graphical tool (Kitematic:https://kitematic.com) on OSX and Windows.

ASaiM is production-ready. It can also be configured to use external accessible computer clusters or cloud environments.

It is also possible and easy to install all or only a subset of tools of the ASaiM framework on existing Galaxy instances. The set of available tools can be easily extended either only a given instance using the Galaxy admin interface or for ASaiM more generally thanks to the simple definition of the installed tools in YAML files available in ASaiM GitHub repository. In the latter case, the Docker image will be automatically rebuilded and the already integrated tools will be updated to keep ASaiM up-to-date. For reproducibility reason, every version of the Docker image is associated to a tag and is conserved.

### Conclusion

ASaiM provides a powerful framework to easily and quickly analyze microbiota data in a reproducible, accessible and transparent way. Built on a Galaxy instance wrapped in a Docker image, ASaiM can be easily deployed with a comprehensive set of tools and their dependencies. These tools are complemented with a set of predefined and tested workflows to address the main microbiota questions (community structure and functions). All these tools and workflows are extensively documented online (http://asaim.readthedocs.io) and supported by Galaxy Interactive Tours and tutorials.

With this complete infrastructure, ASaiM offers a good environment for sophisticated microbiota analyses to scientists without computational knowledge, while promoting transparency, sharing and reproducibility.

## Methods

For the tests, ASaiM was deployed on a computer with Debian GNU/Linux System, 8 cores Intel(R) Xeon(R) at 2.40 GHz and 32 Go of RAM. The workflow has been run on two mock community samples of Human Microbiome Project (HMP), containing a genomic mixture of 22 known microbial strains. The details of comparison analyses are described in the Supplementary Material.

## Availability of supporting source code and requirements

- Project name: ASaiM
- Project home page:https://github.com/ASaiM/framework
- Operating system(s): Platform independent
- Other requirements: Docker
- License: Apache 2

All tools described herein are available in the Galaxy Toolshed (https://toolshed.g2.bx.psu.edu). The Dockerfile to automatically install deploy ASaiM is provided in the GitHub repository and a pre-built Docker image is available at https://quay.io/repository/bebatut/asaim-framework.

## Declarations

### Authors’ contributions

BB, KG, CD, SH, JFB, EP, PP contributed equally to the conceptualization, to the methodology and to the writing process. JFP, PP contributed equally to the funding acquisition. BB, KG, SH contributed equally to the software development and BB, KG, CD and JFP to the validation.

## Acknowledgements

The authors would like to thank EA 4678 CIDAM, UR 454 INRA, M2iSH, LIMOS, CRRI for their involvement in this project, as well as Réjane Beugnot, Thomas Eymard, David Parsons and Björn Grüning for their help.

